# Ancient Darwinian replicators nested within eubacterial genomes

**DOI:** 10.1101/2021.07.10.451892

**Authors:** Frederic Bertels, Paul B. Rainey

## Abstract

Integrative mobile genetic elements (MGEs), such as transposons and insertion sequences, propagate within bacterial genomes, but persistence times in individual lineages are short. For long-term survival, MGEs must continuously invade new hosts by horizontal transfer. Theoretically, MGEs that persist for millions of years in single lineages, and are thus subject to vertical inheritance, should not exist. Here we draw attention to an exception — a class of MGE termed REPIN. REPINs are non-autonomous MGEs whose duplication depends on non-jumping RAYT transposases. Comparisons of REPINs and typical MGEs show that replication rates of REPINs are orders of magnitude lower, REPIN population size fluctuations correlate with changes in available genome space, REPIN conservation depends on RAYT function, and REPIN diversity accumulates within host lineages. These data lead to the hypothesis that REPINs form enduring, beneficial associations with eubacterial chromosomes. Given replicative nesting, our hypothesis predicts conflicts arising from the diverging effects of selection acting simultaneously on REPINs and host genomes. Evidence in support comes from patterns of REPIN abundance and diversity in two distantly related bacterial species. Together this bolsters the conclusion that REPINs are the genetic counterpart of mutualistic endosymbiotic bacteria.

## Introduction

Integrative and replicative mobile genetic elements (MGEs), such as transposons and insertion sequences, are features of both prokaryotic and eukaryotic genomes. While sharing identical mechanisms of replication, the evolutionary fates of MGEs are strongly influenced by differences in genome composition that for the most part distinguish eukaryotes from prokaryotes.

In eukaryotes, where more than half of the genome is non-functional and repetitive (1), duplication of MGEs rarely results in host gene inactivation and thus selection is powerless to prevent expansion of MGE populations (2–5). Continued duplication leads to bloated genomes (6–9) (red (left) area in **Figure 1A**). Unlimited expansion is sometimes counteracted by episodic loss of large parts of the genome (7, 10). Theoretically, stable populations can be achieved through exponential increases in the cost incurred by individual MGEs, or by downregulation of transposition rate (11, 12). In the case of MGEs that contribute fitness benefits to hosts, theory predicts no qualitative change in the fate of MGEs (red (left) area in **Figure 1B**).

**Figure 1.**
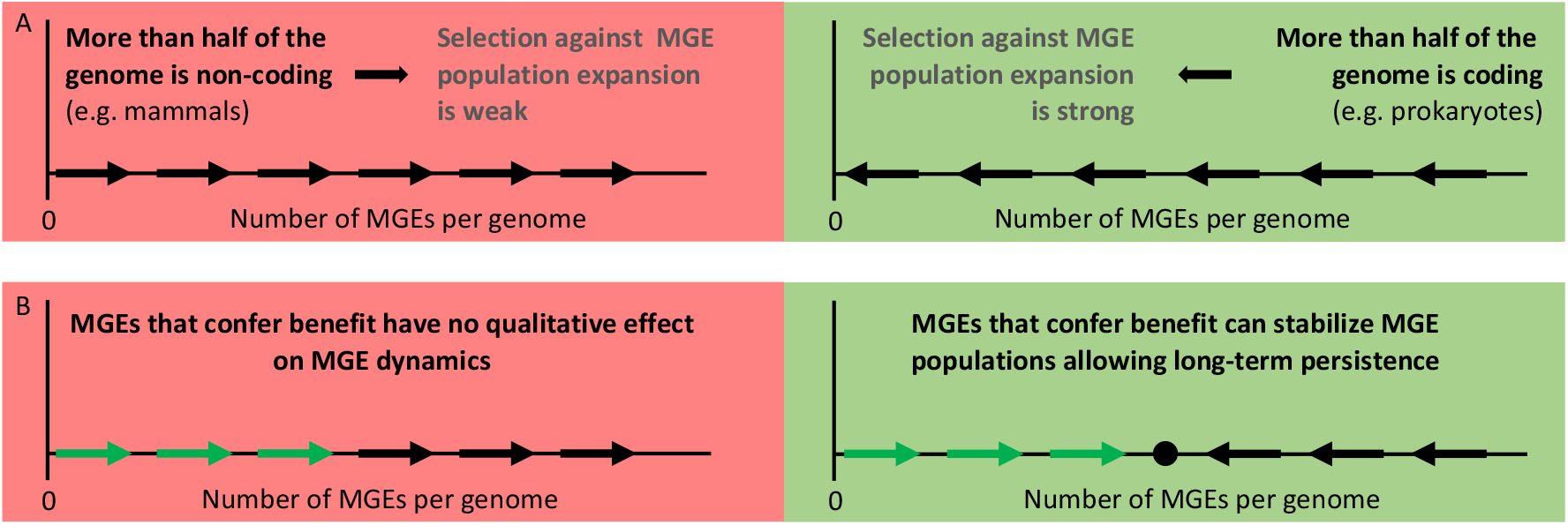
Evolutionary dynamics of MGEs, and dependency on gene content and fitness effects. **(A)** Selection has little opportunity to prevent continual expansion of MGE populations in organisms that harbour large portions of non-coding DNA, for example, eukaryotes, despite fitness costs associated with individual elements (red (left) area). Conversely, in prokaryotes, that contain genomes with little non-coding DNA, selection against MGEs is strong (the majority of duplication events inactive host genes) leading to MGE extinction (green (right) area) (13, 14). **(B)** In the case of MGEs that confer fitness benefits, the presence of such elements in genomes containing substantial portions of non-coding DNA will not alter the drive toward ever increasing MGE population size, although will likely increase the rate of population expansion (red (left) area) (4). However, in genomes with high coding density, presence of MGEs that confer fitness benefits can prevent MGE extinction and lead to long term evolutionary stability (green (right) area). To reach a constant MGE population size the benefit to host must decrease with MGE population size assuming the cost of each MGE for the host is constant (2).

In stark contrast, MGEs are rare in prokaryotes (15). Prokaryotic genomes consist mainly of protein coding genes (gene density > 50%), which means that duplication of MGEs – and concomitant insertion into new genomic regions – stands to inactivate host genes. This has significant fitness costs for both host and MGEs (green (right) area in **Figure 1A**) (2, 3). Consequently, the fate of MGEs in single prokaryotic lineages is extinction, with long term survival being dependent on continual infection (by horizontal transfer) of new hosts (14).

From a theoretical perspective, MGEs can be maintained in prokaryotic lineages over long evolutionary timescales if they contribute some fitness benefit to the host. Long-recognised, are cases of domestication in which a single MGE is co-opted to perform some host-beneficial function, while at the same time, losing ability for autonomous replication (16–18). Notably though, loss of replicative capacity means that such co-opted elements are no longer MGEs. A more intriguing possibility is the existence of persistent, vertically inherited, MGE populations (**Figure 1B, green area**) (2).

Here we outline a case for the existence of such MGEs. The element is a bipartite system involving a **R**EP-**a**ssociated **t**yrosine **t**ransposase (RAYT) and a family of repetitive, short, palindromic, non-autonomous elements termed REPINs (**REP** doublets forming hair**pin**s). RAYTs are incapable of mediating their own replication, but facilitate the replication and persistence of REPINs (18–20).

We begin with a brief description of key features of the REPIN-RAYT system, including likely origin, distribution, persistence, duplication rates and mode of transmission, and show that in all regards these attributes are markedly different from those that define typical bacterial MGEs. We then report analyses demonstrating that REPINs form populations characteristic of living organisms, including evidence of population size fluctuations that correlate with available resources (genome space). Together these data lead to the conclusion that REPINs form enduring, beneficial relationships with eubacterial chromosomes. Moreover, they provoke the hypothesis that REPINs are conceptually similar to beneficial endosymbiotic bacteria that replicate within certain eukaryotic hosts. Given replicative nesting, our hypothesis predicts conflicts arising from the diverging effects of selection acting simultaneously on REPINs and host genomes. Evidence in support comes from patterns of REPIN abundance and diversity in two distantly related bacterial species.

### The REPIN-RAYT system

REPINs are short (∼100 bp long), repetitive (typically 100s of copies per genome), extragenic, palindromic sequences (**Figure 2**) found in approximately 10% of eubacteria. To date, they have not been detected in archaea. REPINs consist of two inverted repeats (REP sequences (21)) each defined by a short (∼14bp) palindrome (green arrows in **Figure 2**) separated by a spacer region (19). The spacer region is highly variable, while the flanking REP sequences are conserved.

**Figure 2.**
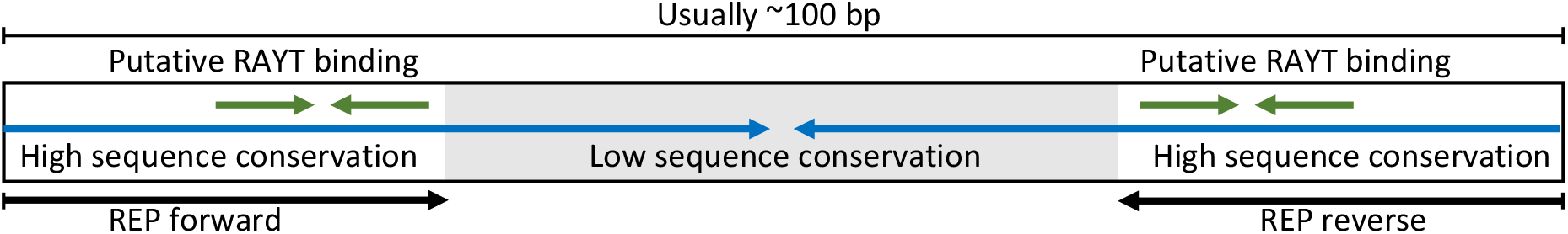
Schematic representation of a REPIN. A typical REPIN consists of two REP sequences in inverted orientation, separated by a short and less conserved DNA sequence (grey highlighting). Blue and green arrows indicate inverted repeats. REPIN length can differ but is usually in the range of about 100bp. REPINs reside almost exclusively in extragenic space.

REPINs are duplicated by a single copy transposase termed RAYT (**R**EP **A**ssociated t**Y**rosine **T**ransposase) (19, 20, 22). Although RAYTs show sequence similarity to transposases, there is no evidence that RAYTs mediate their own transposition. Within bacterial species, RAYTs are typically found in the same extragenic space, are absent from plasmids and other horizontally transferred elements and their phylogeny is congruent with that of the host cell (18). RAYTs are thus vertically inherited and rarely move between bacterial hosts.

RAYTs are highly conserved and originated at least 150 million years ago (based on likely presence of a Group 2 RAYT in the common ancestor of enterobacteria (18)), but are likely much more ancient given the presence of RAYTs across many gammaproteobacterial species. RAYTs probably arose from an IS*200*-like ancestor (see below) since they share identical catalytic domains (HUH and a catalytic tyrosine) and both are associated with short palindromic sequences. Whereas IS*200* sequences form a single cohesive cluster, RAYTs form five sequence groups. All RAYT groups are found as single copy genes in bacterial chromosomes, but only Group 2 and Group 3 RAYTs are associated with REPINs (18).

### REPINs and RAYTs evolved from an IS*200*-like ancestor

The association between both transposases (IS*200* and RAYTs) and short palindromic sequences suggests a common evolutionary origin. IS*200* transposases are flanked by short palindromic repeats that are essential for transposition function (23). While extant IS*200* elements retain a classical transposition-based life history, the lineage that gave rise to RAYTs took a markedly different route: here, palindromic sequences flanking an IS element appear to have fused to form a REPIN capable of exploiting the transposase function for duplication (24). The typical evolutionary fate of such non-autonomous elements is extinction, marked first by loss of full-length transposase genes, followed by degradation of non-autonomous elements (**Figure 3**, (25)). Curiously, RAYT transposases have not been lost. But their inability to replicate independently of the chromosome, means that maintenance of RAYTs must depend on beneficial fitness contributions to the host bacterium. In actual fact, the benefit must arise from a combination of both RAYT and REPIN activity. If this were not so, RAYTs would lose ability to duplicate REPINs, and in turn, REPINs would go extinct.

**Figure 3.**
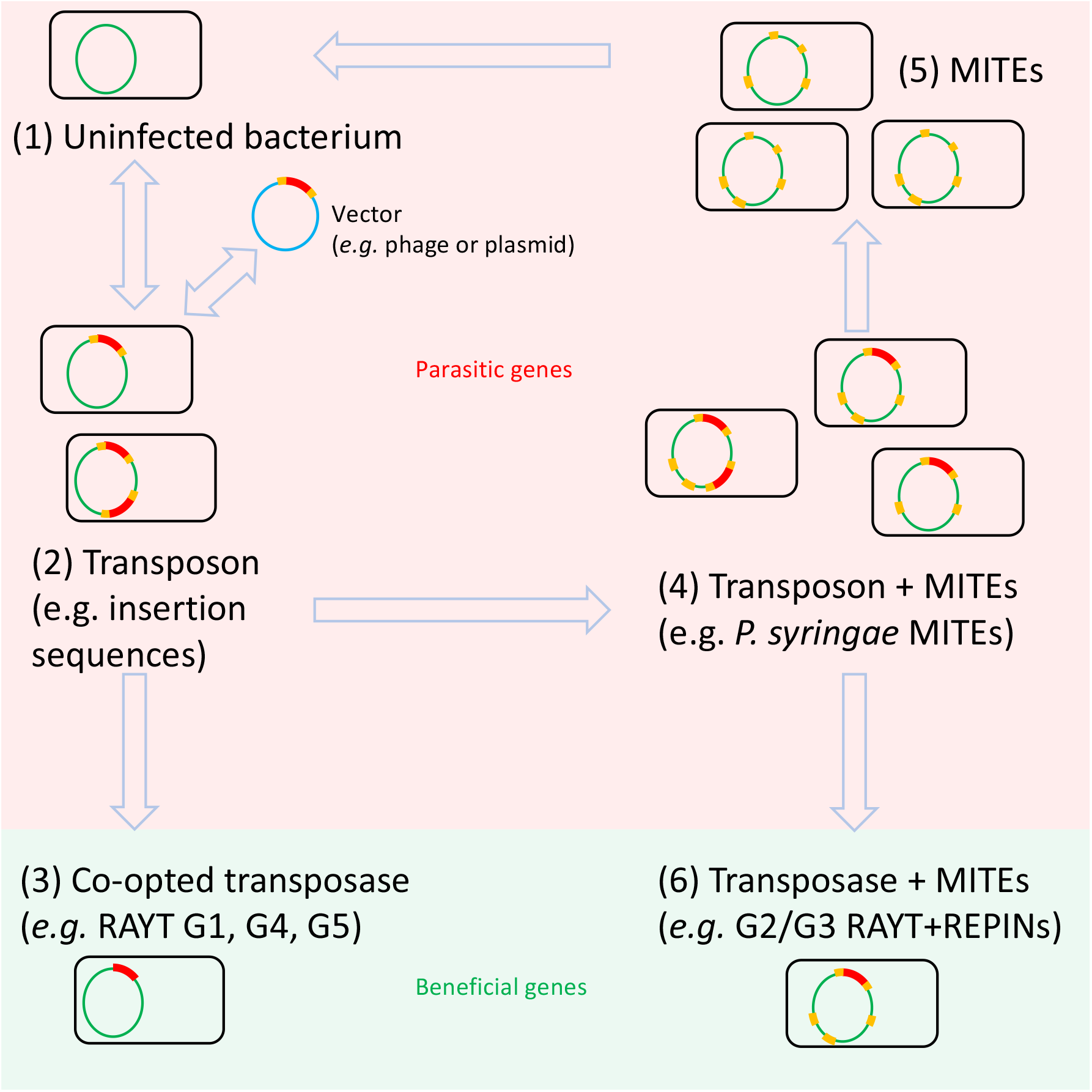
Stages of REPIN-RAYT evolution. (**1**) An uninfected bacterium becomes infected by a vector, such as a plasmid or phage (blue circle), which contains a transposon (for example, an insertion sequence) with flanking sequences (orange) and transposase gene (red). (**2**) The vector infects the bacterium and the transposon inserts into the genome. Once in the genome the transposon may duplicate, go extinct (**1**), or (**3**), on rare occasions, the transposase may be co-opted by the host to perform some host-beneficial function. The latter results in the transposase becoming a single copy gene, and unable to duplicate as observed for Group 1, 4 and 5 RAYTs (18). (**4**) Insertion sequences may also duplicate inside the bacterial genome and leave multiple copies. Sometimes the transposase gene in one of the copies is deactivated. Yet, flanking genes persist and duplicate by parasitizing transposase function encoded elsewhere in the genome. Flanking sequences lacking transposase function are called miniature inverted repeat transposable elements (MITEs) (26). (**5**) MITEs can parasitize full-length transposons and may even cause loss of transposon movement. Once the last functional transposase is eradicated from the genome, MITEs face extinction (25, 28). (**6**) Alternatively, both transposases and associated MITEs are co-opted by the host bacterium. This is the case for Group 2 and Group 3 RAYTs and their associated REPINs. REPINs differ markedly from MITEs. While REPINs duplicate slowly (∼10^−8^ per host replication) and persist for long periods of time (29) MITEs duplicate as fast as their parental elements (∼10^−5^ (30)) and persist only briefly before extinction.

By their nature, REPINs share many similarities with **m**iniature-**i**nverted repeat **t**ransposable **e**lements (MITEs) (26, 27), which like REPINs, have also evolved from repeat sequences flanking transposons (**Figure 3**). REPINs, however, are distinct from MITEs in several important regards. Firstly, the evolution of MITEs occurs repeatedly – and from diverse transposon types – with short persistence times that are tied to the fate of the cognate transposon. In contrast, REPIN-RAYT systems have evolved rarely – possibly only once – and have persisted for millions of years (see below and (18)).

Secondly, and a primary distinguishing feature, is the nature of the interaction between MITEs and their cognate transposases, and REPINs and their cognate RAYTs. MITEs directly parasitise transposon-encoded transposase function – with presumably detrimental effects on transposon fitness – whereas parasitism is not possible in the case of REPINs and their cognate RAYTs. This is because RAYT-encoded function does not mediate RAYT duplication. While the relationship between REPIN and RAYT most likely began as exploitative it appears to have evolved toward a beneficial association (see below) – both in terms of the interaction between RAYT and host cell, and between RAYT and REPIN (2, 19).

Thirdly, and a likely consequence of the second point, is duplication rate. MITEs duplicate about once every 100,000 host-cell generations (30), whereas REPINs duplicate just once every 100 million host-cell generations (see below and (29)). A faster rate of duplication is expected in the case of an exploitative interaction, with selection likely to favour ever increasing rates of transposition until the point of either host-cell extinction, or extinction of the cognate transposon (2, 31).

### REPIN sequence diversity is evidence of long-term persistence

REPINs persist over millions of years and display patterns of molecular evolution distinct from MGEs such as transposons and insertion sequences. Evidence comes from analyses of REPIN diversity. Central to such analyses is the relationship between the rate of duplication and the number of mutations per sequence replication (per site mutation rate multiplied by sequence length). Elements whose duplication rate is much higher than the mutation rate show little within-element diversity, whereas diversity accumulates (through mutational decay) in elements that duplicate at rates similar to the mutation rate. These mutation dynamics are described by Quasispecies theory (also known as the mutation-selection model (32)). According to Quasispecies theory, the most common REPIN sequence in the population defines the “master sequence” (mutation class 0 in **Figure 4**). Other sequence categories are defined by the number of nucleotides different from the master sequence. For a given mutation rate the model predicts that REPIN duplication rates are in the order of 10^−8^ duplications per host generation (29). This means that REPINs are duplicated at a rate that is at least three magnitudes slower than duplication rates of insertion sequences (13).

**Figure 4.**
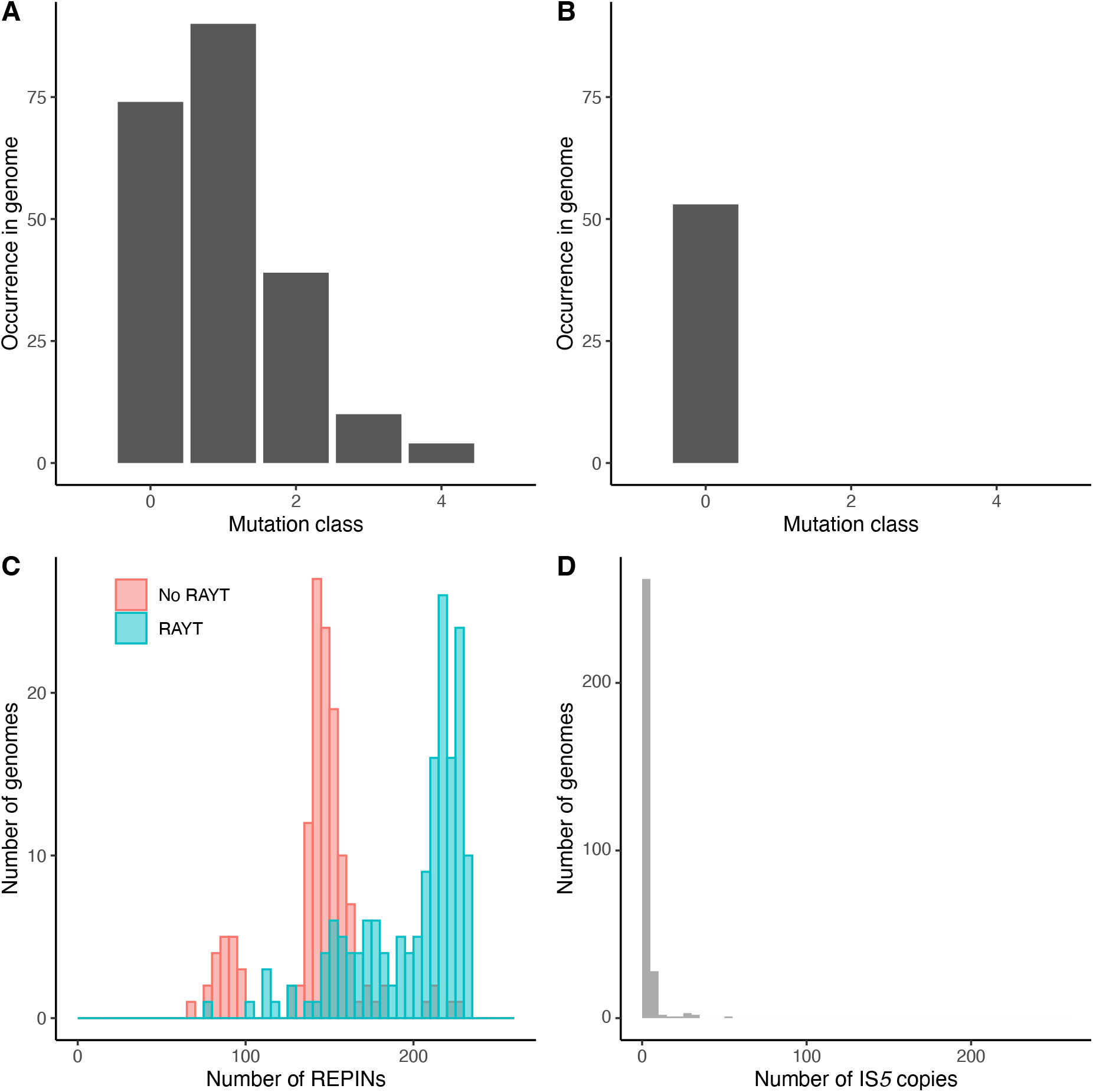
REPIN sequence diversity within genomes and population size distribution across genomes are indicative long persistence times. (**A**) Sequence distribution of the most frequent 21 bp long sequence (part of a REPIN) as well as all relatives from *E. coli* L103-2. Mutation classes are a Quasispecies concept. Mutation class 0 is the master sequence, the most common sequence in a population. Mutation class 1 contains all sequences that differ from the master sequence by exactly one nucleotide. Mutation class 2 are all sequences in the population that differ by exactly two nucleotides and so on. (**B**) The same data as in (A) for the most common 21bp sequence found as part of IS*5* in *E. coli* L103-2. This is also the *E. coli* strain that contains the most IS*5* copies across all 300 strains analyzed. (**C**) REPIN population size across 300 *E. coli* strains (duplicate genomes are not included). Note that *E. coli* strains not containing RAYT genes contain significantly fewer REPINs. (**D**) IS*5* copy numbers in the same 300 *E. coli* genomes. IS*5* copy numbers are skewed to the left of the graph, whereas REPIN population sizes are more reminiscent of a normal distribution with a mean of about 143 (mode of 142) in the case where no RAYT genes are present in the genome (red) or about 200 (mode of 217) where RAYTs are present in the genome (turquoise). (C and D) adapted from Park et al. (2).

Low REPIN duplication rates are consistent with long REPIN persistence times. A REPIN must be present within a genome for approximately 100 million host-cell generations in order to duplicate. During this time, REPINs accumulate on average about 0.2 mutations. Hence, for a REPIN to acquire four mutations it must be maintained inside a host genome for approximately 800 million generations. Accumulation of diversity within REPINs thus requires very long REPIN persistence times.

High levels of REPIN sequence diversity are evident in *E. coli* (29). For example, in *E. coli* L103-2, less than half of all REPINs match the master sequence category. The remainder show a degree of divergence from the master sequence due to acquisition of mutations over extended time periods (**Figure 4A**). Yet, across the entire *E. coli* species the master sequence has not changed: it is identical in every strain.

By way of contrast, the exact same analysis for the insertion sequence IS*5* in *E. coli* L103-2, shows no diversity across IS*5* copies within a single genome (**Figure 4B**). Absence of IS sequence diversity has been observed previously (33), and demonstrates that insertion sequences do not persist long enough to diversify within genomes, as has been observed previously (33, 34).

Short persistence times of insertion sequences and long persistence times of REPINs can also be observed in comparisons made among genomes. Across 300 *E. coli* genomes that encompass the currently known sequence diversity of *E. coli*, there is not a single genome from which REPINs are absent (**Figure 4C**). This means that REPINs have been maintained within *E. coli* for at least the last 15 million years, ever since divergence from the most recent common ancestor (29). Interestingly, some *E. coli* strains have lost the RAYT gene and REPIN population sizes in such RAYT-less genomes is significantly reduced, consistent with the role that RAYTs play in maintenance of REPIN populations. In contrast to REPINs, IS*5* is present in less than half of all *E. coli* genomes (133 out of 300, **Figure 4D**). The patchy presence of insertion sequences has been previously reported and indicates, together with the lack of genetic variation within genomes, that insertion sequences are frequently purged from genomes, with constant reinfection of new hosts being necessary for persistence (13).

### REPINs are not parasites

In a recent theoretical study factors affecting persistence of REPINs have been explored (2). According to the developed model, REPINs are assumed to confer no fitness costs to hosts, however, costs arise on transposition because REPIN movement stands to inactivate genes necessary for host survival. The probability that a host lineage dies after such a transposition event is set by a parameter termed gamma. Gamma determines the proportion of the genome that is functionally important, namely, is required for survival in environments that the host is likely to encounter over the long term. In *E. coli*, for example, these are likely to be genes that are conserved across the species (35).

In the presence of horizontal transfer, equilibrium of the system is entirely determined by the value of gamma. If more than half of the genome encodes for functionally important genes (gamma > 0.5) then the fate of MGEs (including REPINs) is extinction. This is because each duplication causes a decline in the MGE population as a consequence of host death. The model accurately predicts the inability of MGEs to persist in prokaryotes (gamma > 0.5), and population expansion in eukaryotes (gamma < 0.5) (10, 34). However, the model fails to explain the persistence of REPINs.

To account for persistence of REPINs it is necessary to assume a fitness benefit to the host that decreases with REPIN population size to counteract a constant cost per REPIN. If REPINs confer some host benefit then REPINs form stable and persistent sequence populations. If the benefit function is removed, such that REPINs are either neutral or parasitic, then REPINs go extinct. This outcome is independent of whether REPINs parasitize RAYTs, or the host (or both). If RAYTs are parasitized by REPINs and REPINs are inadvertently duplicated by RAYTs, then the fate of REPINs is extinction. Extinction is assured for the reasons given above and irrespective of duplication rate: every time a REPIN duplicates the number of REPINs residing in a bacterial population decreases because REPIN duplication tends to kill the host bacterium.

### REPIN populations respond to available niche space

REPINs vary one to another, they replicate and leave offspring copies that resemble parental types. They are thus Darwinian entities in their own right, replete with their own population biology that can be read from patterns of diversity and abundance within bacterial genomes. As with any population of entities, population size depends on available resources. For example, 1 mL of rich broth medium allows *E. coli* populations to reach ∼10^9^ individuals; 1 L allows the population to expand to 10^12^ cells. In both volumes the density of cells at stationary phase remains the same. REPINs are expected to behave similarly – and drawing a parallel between genome size and resource availability – a REPIN population of 100 individuals in a genome of 1 Mb, should increase 10-fold with a 10-fold expansion in genome size. REPIN density should remain constant.

As shown in **Figure 5**, across 2828 representative bacterial chromosomes, the density of 16-mer repeats (a proxy for REPINs abundance in genomes that contain a RAYT gene) remains constant for genomes that contain Group 2 or Group 3 RAYT genes (only Group 2 and Group 3 RAYTs are associated with REPINs (18)). Repeat density decreases with increasing genome length in the absence of RAYT genes. Even in genomes that contain IS*200* genes, the most closely related IS transposase, repeat density decreases with increasing genome length. These data indicate that REPIN population size does indeed increase with available niche space, exactly as expected for populations of any biological entity.

**Figure 5.**
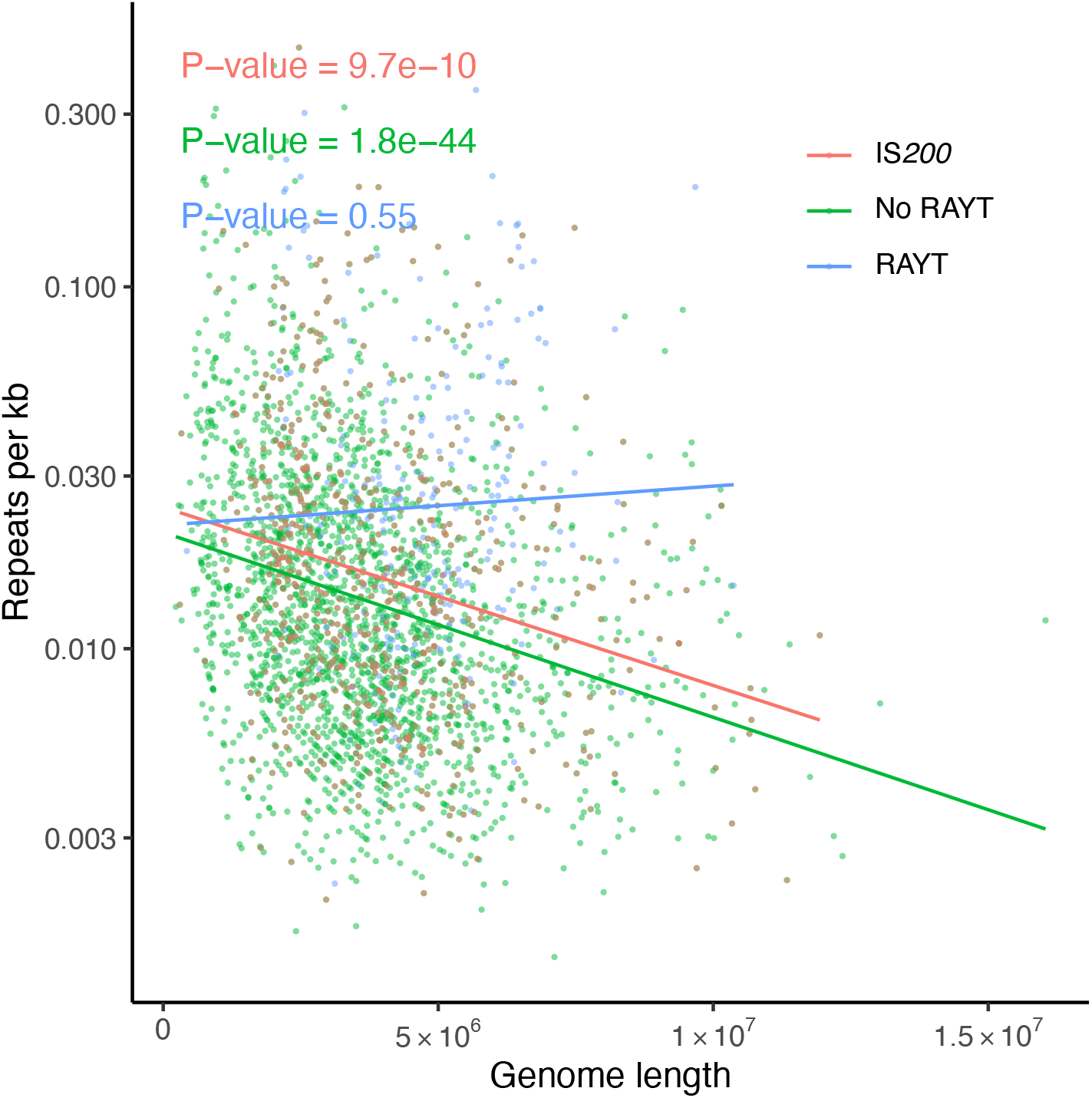
Repeat density remains constant in genomes containing RAYT genes. Repeat density (the frequency of the most abundant 16 bp sequence divided by genome length) significantly decreases in chromosomes that do not contain RAYT genes (2567 chromosomes, green) as well as chromosomes that do not contain RAYT genes, but do contain IS*200* genes (596 chromosomes, orange). Repeat density does not significantly decrease in genomes containing either Group 2 or Group 3 RAYTs (232 chromosomes, blue). The differences in p-Values cannot be attributed to sample size alone: when sampling 232 genomes from 2567 genomes 1000 times, all samples showed a negative slope in contrast to the positive slope observed for RAYT-containing genomes. In 992 of 1000 samples the slope is significantly different from 0, with a p-Value of less than 0.05. Data are derived from the analysis of 2828 representative bacterial chromosomes downloaded from NCBI on the 16.11.2020 https://www.ncbi.nlm.nih.gov/genome/browse/.

Similar to natural populations of, for example, plants or animals, that grow in a given environment, size of that environment is not the only parameter that determines carrying capacity. Nutrient content, predators and other factors will exert a strong effect on carrying capacity. Similarly, high variance in repeat density in **Figure 5** suggests factors other than genome size play major roles in determining REPIN population size in bacterial genomes.

### REPINs resemble beneficial endosymbiotic bacteria

Conceptually, REPINs resemble beneficial endosymbiotic bacteria. REPINs are non-infectious, non-autonomous entities that replicate within bacterial chromosomes. They are distinct from other replicative sequences, such as transposons, insertion sequences, plasmids and integrative and conjugative elements, in that they are transmitted vertically, they are not parasites, they form stable, enduring populations, and have persisted across entire species for millions of years (**Figure 6**).

**Figure 6.**
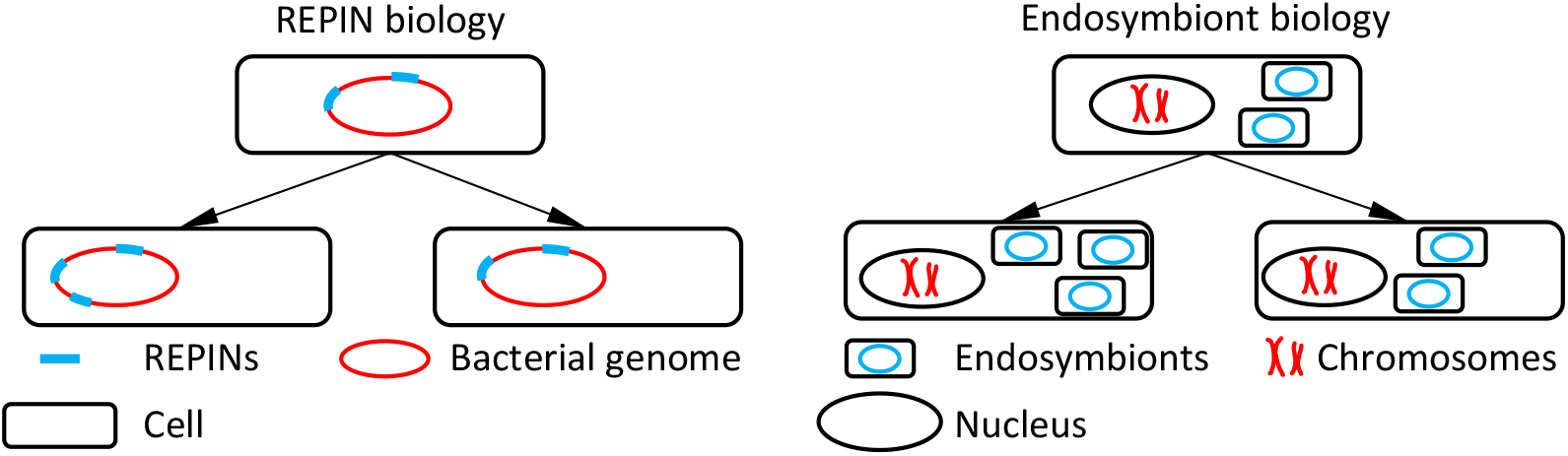
REPIN biology resembles endosymbiont biology. The left panel depicts REPINs that reside in genomes, that duplicate and are passed on vertically to bacterial offspring. REPIN duplication inside the chromosome is mediated through the RAYT transposase at extremely low rates. The right panel shows endosymbionts residing in the cells of an organism as for example, bacteria, such as *Rickettsia* inside arthropod cells, or mitochondria inside the eukaryotic cell. Here, endosymbionts duplicate and are passed on vertically to offspring. Chromosomes of endosymbionts are replicated independent of host chromosomes, while REPINs are replicated as part of the host chromosome.

The association between REPINs and host genomes is facultative – at least on part of the host – thus bringing the resemblance of REPINs closer to facultative endosymbionts, such as *Hamiltonella* and whiteflies, than to obligate endosymbionts, such as *Buchnera* and aphids, or mitochondria and the eukaryotic cell (36). RAYTs can be deleted from bacterial genomes, thus depriving REPINs of replicative capacity and eliminating presumed benefits to host, without noticeable effects on cell growth (37). Moreover, through evolutionary time, RAYTs have been lost from host genomes accompanied by gradual erosion and loss of REPIN populations (**Figure 7A**). Despite loss, descendant lineages remain viable.

**Figure 7.**
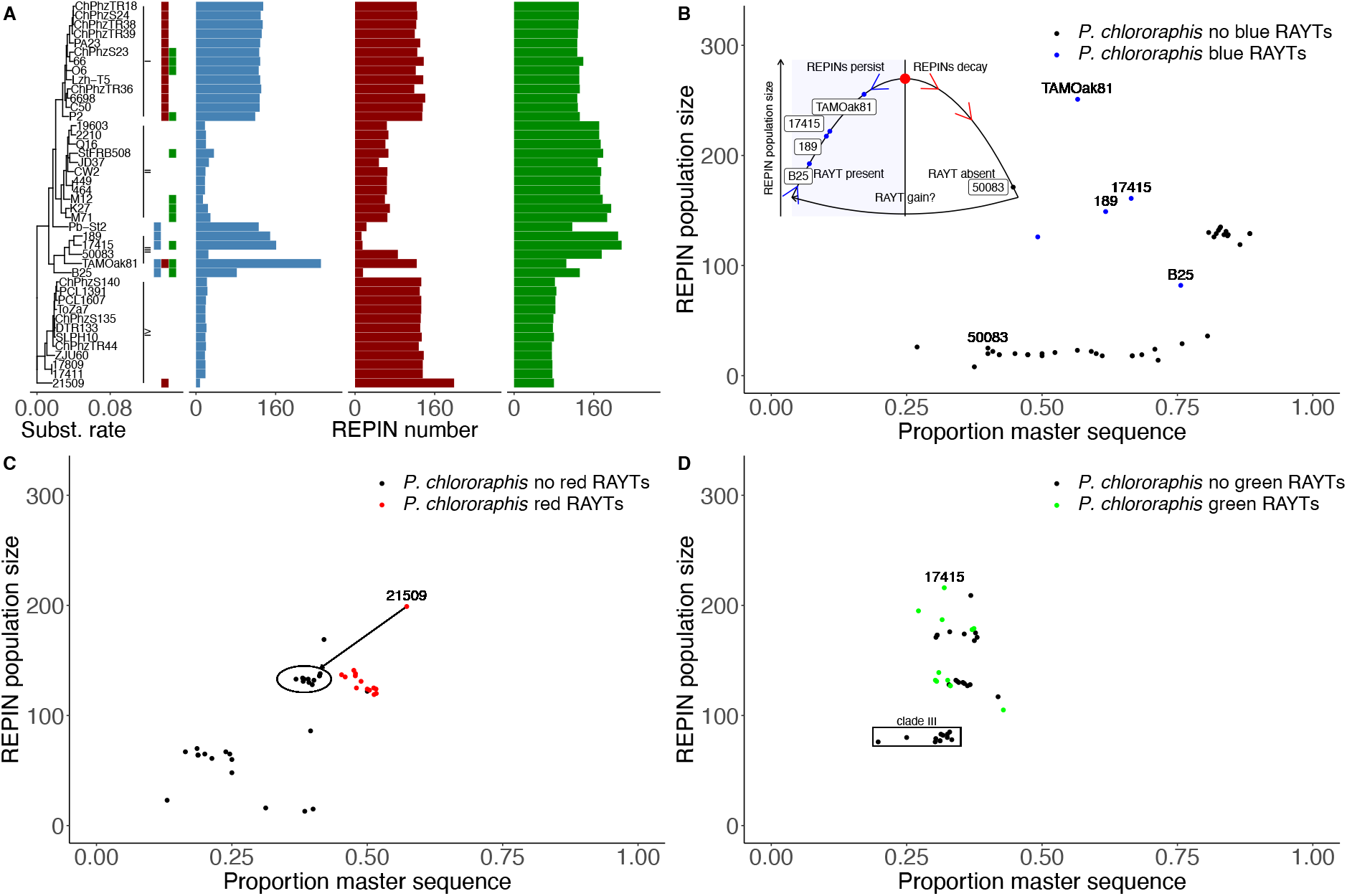
RAYT presence and absence determines REPIN dynamics in *Pseudomonas chlororaphis*. (**A**) The tree on the left side of the figure shows a phylogeny of different *P. chlororaphis* strains. The tree was built based on a whole genome alignment generated via REALPHY (44). REALPHY was run on the webserver (version 1.13) using TAMOak and 17411 as references. The resulting alignments were merged resulting in a total of 3,850,960 alignment positions. From this alignment a phylogeny was generated with a GTR substitution model using PhyML (45). The same tree is shown with bootstrap values in **Supplementary Figure 2**. The next column shows the presence and absence of three different RAYT transposases (blue, red and green). (**B-D**) The proportion of master sequences (indicates sequence conservation) in a REPIN population and the REPIN population size. According to Quasispecies theory or mutation-selection balance, the higher proportion of master sequences (the most common sequence in the population) correlates with higher duplication rates of the sequence population. The inset in (B) shows a plausible cyclical dynamic of REPINs arising from the tension between REPIN and cell-level selection. The blue arrows in the inset indicate a stable evolutionary state of a REPIN population in the presence of a corresponding RAYT gene. A second evolutionary stable state is indicated by the red arrows in the absence of the RAYT gene. When RAYT genes are absent REPINs decay and eventually vanish from the genome. A REPIN population could in theory be rescued through the acquisition of a corresponding RAYT gene. See text for details.

However, unlike facultative endosymbiotic bacteria that are transmitted horizontally, can switch hosts (38, 39) and live independently of hosts (36), REPINs are almost exclusively passed to offspring genomes by vertical transmission. While horizontal transfer is possible, establishment of a REPIN population in a new host requires that there be a resident RAYT with which the transferred REPIN is compatible. Given the specificity between RAYTs and REPINs within individual lineages – a reflection of persistent co-evolution between REPINs and their cognate RAYT ((19) and **Figure 4**) – the chances of transfer leading to establishment of a new REPIN population is vanishingly small. A related issue is the complete absence of RAYTs from plasmids and other horizontally mobile agents suggesting that carriage of RAYTs on such elements is costly or simply not beneficial to the plasmid (18, 40).

An additional departure of REPINs from the life history of facultative endosymbionts stems from semiconservative replication of the bacterial genome. Whereas facultative endosymbionts often recolonise hosts from limiting numbers of cells, that is, they frequently passage through population bottlenecks, chromosome replication during cell division means each daughter cell receives equal numbers of REPINs. In this regard, REPINs have much less autonomy compared to even obligate, vertically transmitted, endosymbionts: each generation of an endosymbiont host requires numerous divisions of the endosymbiont, whereas REPINs replicate just once in about 100 million host generations.

REPINs also resemble obligate endosymbionts, in terms of their autonomy. For replication, REPINs are dependent on a single copy RAYT gene encoded by the host genome. As shown in **Figure 4**, in the absence of the RAYT gene, REPIN populations slowly decay. Obligate endosymbionts also rely on host encoded functions for replication (36). For example, the DNA of mitochondria is replicated by polymerase gamma, a host encoded protein. Apart from polymerase gamma there are over 1000 other mitochondrial proteins that are encoded in the nuclear DNA (41).

A related issue concerns the consequences of replication. Replication of endosymbionts is essential for re-establishment of functional (beneficial) interactions with hosts. While excessive replication could potentially cause harm, for the most part this is not possible, because endosymbionts are often constrained to particular organismal structures, such as, in the vibrio-squid symbiosis, crypts of the bobtail squid (42). In the case of REPINs, duplication is likely to be costly.

### Signatures of intragenomic conflict

Interactions among independently replicating Darwinian populations, such as those that exist between endosymbiotic bacteria and their hosts, or mitochondria and the eukaryotic nucleus, establish conditions that render conflict likely. In instances where one Darwinian population is nested within another, conflict is assured (43). If our hypothesis that REPINs are conceptually similar to endosymbiotic bacteria is correct, then selection at the level of REPINs is predicted to favour variants with enhanced replicative capacity, even though heightened REPIN activity stands to harm host cell lineages. In turn, cell-level selection is expected to reduce REPIN replication to a level that minimizes harm, most potently *via* effects on RAYTs. While, over evolutionary time, selection will favour those cell lineages containing REPINs whose replicative capacity most closely aligns with the long-term fate of cells, co-existence is likely to be punctuated by periods of conflict.

Ability to capture evidence of conflict – and possibly also conflict resolution – depends on when in the past conflict occurred, the time scale and mode of resolution, and whether signatures of conflict are sufficiently conserved in extant lineages. The inset to **Figure 7B** shows a plausible cyclical dynamic. The blue trajectory on the left depicts the zone of REPIN persistence. Increases in REPIN population size are driven by REPIN-level selection, but counterbalanced by cell-level selection. While persistence within the blue zone is expected to benefit both host and symbiont, REPIN-level selection may nonetheless drive REPIN numbers beyond a critical threshold to the point where cell-level costs become unsustainable. Such instances will lead to extinction of affected cell lineages, or elimination of RAYT function. The latter will result in REPIN decay (both in number and match to the master sequence) and is indicated by the red trajectory. Decay is expected to be a gradual process unfolding over the course of millions of years. There remains the possibility of recovery if RAYT function can be restored.

We turn to two sets of closely related genomes in which unusual patterns of REPIN-RAYT evolution are suggestive of long-term evolutionary conflicts. The first is a set of 42 strains of the plant associated bacterium *Pseudomonas chlororaphis*; the second is a set of 130 strains of *Neisseria* including *N. gonorrhea* and *N. meningitidis*. Both lineages display similar levels of sequence divergence (about 0.04 substitutions per site), similar also to the divergence observed in *E. coli* (29, 46). If we assume that evolutionary rates are comparable between the two lineages and E. coli, then the *P. chlororaphis* species diverged approximately 15 million years ago; the same holds for the two *Neisseria* species.

**Figure 7A** shows the phylogenetic relationship among *P. chlororaphis* strains. The coloured (blue, red and green) boxes indicate the presence / absence of three different RAYT transposases. Each RAYT type is found at the same position in the *P. chlororaphis* genome and forms a monophyletic group in the RAYT phylogeny (**Supplementary Figure 1**). It is likely that the ancestral genotype contained all three RAYTs. Also depicted in **Figure 7A** is the REPIN population size corresponding to each RAYT type.

Patterns of REPIN evolution most consistent with conflict and resolution can be seen in clades III and IV. Focusing firstly on clade III the ancestral type of this set of five strains contained a blue RAYT. Strains B25, 189, 17415 and TAMOak81 retain the blue RAYT: each harbours between 80 and 250 REPINs (blue labelled dots in **Figure 7B**, also indicated on the inset) and more than 60% of these REPINs are an exact match to the master sequence. This is in striking contrast to 50083 (black labelled dot in **Figure 7B** and also inset) from which the blue RAYT has been eliminated. Strain 50083 contains few REPINs with few matching the master sequence.

Clade IV comprises 12 strains that, with the exception of strain 21509, have lost all three RAYTs. Based on patterns of REPIN abundance and proportion of master sequence it is likely that blue RAYTs were the first to be eliminated, followed by green and lastly red. However, 21509 has retained the red RAYT, with this strain containing more than 200 associated REPINs and 60% matching the master sequence (**Figure 7C**). In the 11 strains lacking the red RAYT the number of REPINs has declined along with matches to the master sequence (black circled dots in **Figure 7C**).

Turning to the green RAYT family, patterns of conflict are less evident than for blue and red, with green RAYTs being evident in three of the four marked clades. This suggests that loss of green may be overall more recent and thus signatures of conflict less prominent. In clade IV, where green RAYTs are absent, decay of REPINs is apparent (black rectangle in **Figure 7D**). It is notable that 17415 (Clade III), which contains a green RAYT, harbours the largest number of REPINs. The sister taxon (strain 189) has lost the green RAYT, possibly recently, and contains the second largest REPIN population.

An additional observation warranting comment concerns clade I. This clade harbours large numbers of all three REPIN families. The red RAYT is present in all strains, with the green RAYT evident in just four strains, while blue RAYTs are absent. Lack of blue RAYTs begs an explanation for the abundance of blue REPINs. A distinct possibility is that red RAYTs have evolved the capacity to aid persistence of both red and blue REPINs. Such a possibility is consistent with the fact that the master sequences of blue and red REPINs differ by just four nucleotides over the 42 nucleotides that define the master sequence. This observation suggests that RAYTs can evolve to maintain and even duplicate closely related REPIN populations, when the cognate RAYT is lost. How a RAYT host switch might affect the sequence identity of a REPIN population is unclear, but is an interesting question for future research.

**Figure 8A** shows the phylogenetic relationship among strains belonging to two different *Neisseria* species, the number of RAYTs, and corresponding REPIN population size. Both species contain the same RAYT type (Group 2 RAYT) and all REPINs share the same identical master sequence. Strains of the *N. gonorrhoeae* clade all share a non-functional RAYT caused by a frameshift mutation (red bars in **Figure 8A**), and all show evidence of REPIN decay (blue dots in **Figure 8B**). In contrast, most strains of *N. meningitidis* harbour functional RAYTs and show no sign of REPIN decay (green dots in **Figure 8B**).

**Figure 8.**
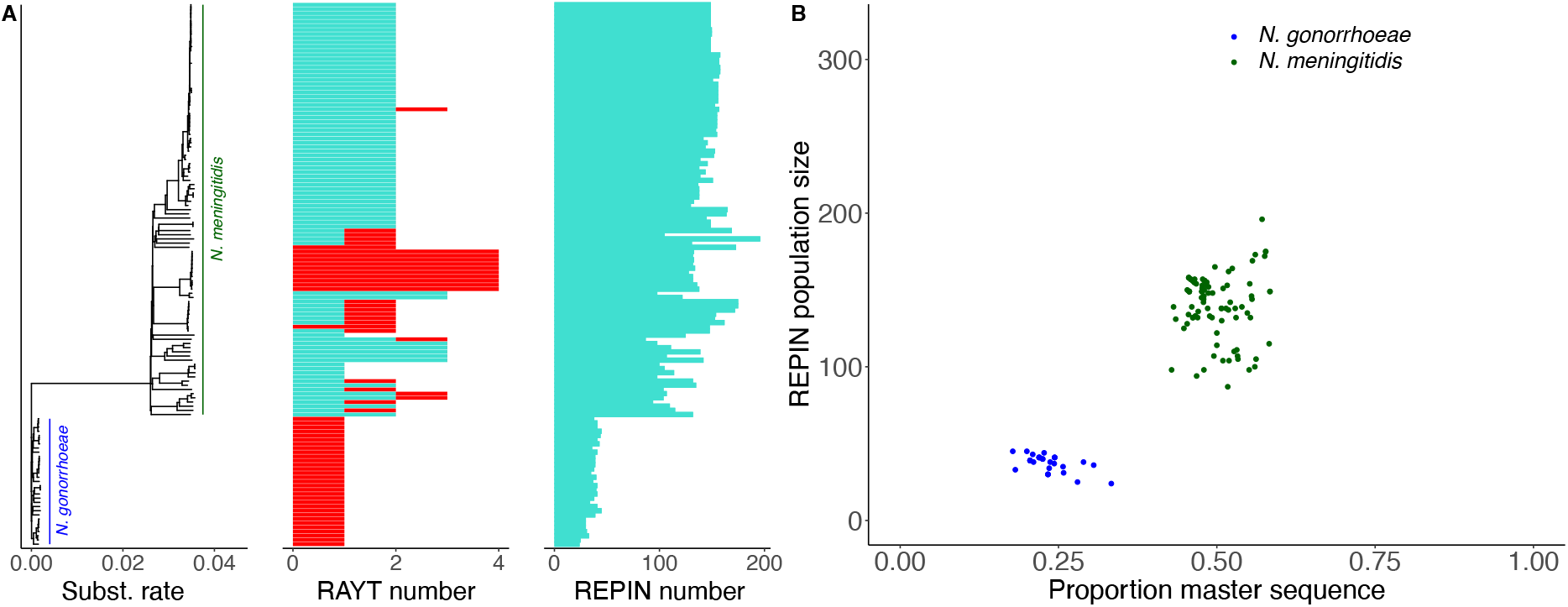
REPIN population size is regulated in *Neisseria*. **(A)** The tree on the left side of the figure shows a phylogeny of different *N. meningitidis* and *N. gonorrhoeae* strains. The first barplot shows the number of RAYT transposases found in each strain. The second barplot shows the number of REPINs present in the largest sequence cluster. The tree was built by applying neighbor joining (47) to a distance matrix generated with “andi” from complete genomes (48). RAYT genes highlighted in red contain frameshifts that lead to a premature stop codon (**Supplementary Figure 3**), **(B)** The proportion of master sequences (proxy for REPIN duplication rate and sequence conservation) for REPIN populations found in *N. gonorrhoeae* (blue) and *N. meningitidis* (green) plotted in relation to population size. Once the RAYT is lost REPIN sequences decay and populations shrink (moving to the bottom left of the graph), as observed for populations from *N. gonorrhoeae*.

A curious feature of *N. meningitidis* is the presence of multiple RAYT copies. This is likely caused by *in trans* transposition, because all RAYT genes are linked to an IS*1106* fragment. Transposition is likely mediated by a full length IS*1106* copy that is present in multiple copies in *N. meningitidis* but absent from *N. gonorrhoeae*. While some RAYTs are present in multiple copies, not all of these are functional. Intriguingly, all non-functional RAYTs in *N. meningitidis* – as in *N. gonorrhoeae* – are caused by frameshift mutations (red bars in **Figure 8A**). In most instances the frameshifts occur in short homopolymeric tracts (**Supplementary Figure 3**) raising the possibility that gain and loss of RAYT function *via* replication slippage may be an adaptive mechanism allowing tension between interests of REPINs and cells to be readily resolved (49).

The dN/dS value of *Neisseria* RAYTs, from all pairwise comparisons (including non-functional copies that contain stop codons) is 0.74. If non-functional RAYT copies are excluded, the dN/dS is much lower (0.4) in line with expectations of increased purifying selection. In *P. chlororaphis* genomes, where only a single functional RAYT is present the dN/dS value is even lower at 0.2. The higher value for *Neisseria* RAYTs probably reflects relaxed selection due both to non-functional copies, but also the presence of up to four functional copies per genome.

While presenting the *Neisseria* data set in the context of conflict between the interests of REPINs and host cells, an alternate possibility is that the change in niche preference associated with divergence of *N. gonorrhoeae* from *N. meningitidis* led to selection against functional REPIN-RAYT systems in *N. gonorrhoeae*. Distinguishing between these competing hypotheses is not possible on the basis of current data, but the apparent maladaptive nature of REPIN-RAYT systems in *N. gonorrhoeae* may warrant future investigation. With reference to the inset of **Figure 7B**, this would be the equivalent of changes in the environment leading to alteration in the balance of effects arising from selection on REPINs versus selection at the level of cells – in effect, a “squashing” of the triangle.

## Conclusion

REPINs are non-autonomous replicative entities (MGEs), that in conjunction with cognate RAYTs, form enduring endosymbiont-like relationships with eubacterial chromosomes.

REPINs form vertically transmitted sequence populations that have persisted within single lineages for hundreds of millions of years, have population biology typical of cellular organisms and, as expected – given nesting within higher order structures – show evidence of periodic tension between the replicative interests of REPINs and host cells. Given our current understanding of how biology works, these properties and features mean that REPINs (plus RAYTs) must provide benefit to host cells (50).

The precise nature of the contribution that REPIN-RAYTs make to host cell fitness is unclear, but studies of REP function over many years reveal contributions ranging from localised effects on mutation and recombination rate (51), to facilitation of genome amplification (52, 53), and modulation of tertiary genome structure (54–56). REPINs can act as preferential targets for insertion sequences (57, 58), or have specific effects on gene expression, either through transcription termination (59), or effects on the stability of mRNA (60–62). Thus far, there have been no studies that link REPIN function to that of RAYT endonucleases, with the exception of recent work showing that RAYTs recognise and cleave single stranded REP DNA (22, 63). Functional studies are underway, with mounting support that the REPIN-RAYT system plays a role in modulation of mRNA stability under conditions of environmental change.

## Materials and Methods

Graphs were generated in R using the packages ape, ggplot2, ggtree and cowplot.

### Identifying REPINs in bacterial genomes

REP and REPIN sequences were identified within bacterial species using a specific program that is available through a webserver (http://rarefan.evolbio.mpg.de) (64).

REP sequence groups were identified using a previously described procedure (19). First, we extracted all 21bp long sequences that occurred more than 55 times in the genome (*P. chlororaphis* TAMOak81, and *Neisseria meningitidis* 2594). We then grouped all sequences that occurred in the genome at least once at a distance of less than 30bp.

For *Neisseria* this procedure identified six REP sequence groups (65). Group 1 and Group 6 yielded identical REPINs (most common 21bp long sequence: ATTCCCGCGCAGGCGGGAATC occurs 241 times in *N. meningitidis* 2594). None of the remaining four sequence groups yielded REPINs (i.e., they did not occur in opposite orientation at a distance of less than 130bp). Group 2 (CTTGTCCTGATTTTTGTTAAT occurs 207 times) overlaps with the CORREIA sequence TATAGTGGATTAACAAAAATCAGGACAA (66, 67).

There are six REP sequence groups in *P. chlororaphis* (68). Group 1 and Group 6 generated the same REPINs, so did Group 2 and Group 4 as well as Group 3 and Group 5. We analyzed REPINs formed by REP Groups 1 to 3, which we assigned the colours blue, red and green respectively. We did not include Group 6 in our analysis, because the number of REPINs contained was low (<40) and the REPINs were not associated with a RAYT transposase. Each REP sequence group was uniquely defined by the most common 21bp long sequence in that group. The most common 21bp long sequences in the first three sequence groups were: TAGGAGCGAGCTTGCTCGCGA (blue, occurs 466 times in TAMOak81); TCGCGGGCAAGCCTCGCTCCT (red, occurs 238 times); CGCAGCCTGCGGCAGCGGCTA (green, occurs 226 times).

From these REP sequence groups, we determined REPINs (two REP sequences in inverted orientation) across all strains in this study as described previously (29). First, from the REP seed sequence we generate all single point mutations. We then determined whether any of the generated mutant 21 bp long sequences are present in the genome. If so, we recursively generated all possible point mutants of the sequences present in the genome until no more sequences were found (files ending in “.ss.REP” (65, 68)). For all identified REP sequences genome positions were determined. Any two sequences that occur at a distance of less than 130bp, with the two sequences occurring on opposite strands (in opposite orientations), are defined as forming a REPIN. Although an entire REPIN consists of three parts (conserved 3’ and 5’ sequence as well as the variable spacer region), here we only analyzed evolution of the 3’ and 5’ conserved regions of the REPIN (found in files ending in “.ss” (65, 68)). The files contain fasta formatted sequences where the sequence name contains all the genomic positions, at which the REPIN was identified. The sequence is a concatenation of the 5’ and 3’ 21bp sequence of the REPIN. REP singlets are all sequences, for which a partner sequence could not be identified. In this case 21As are concatenated to the end of the sequence.

Files are categorized into subfolders starting with the strain name and ending in “_[number]”, where the number corresponds to the REPIN type. There is only a single REPIN type in *Neisseria* (i.e. ending in “_0”). In *P. chlororaphis* there are three REPIN types: blue ending in “_0”, red ending in “_1” and green ending in “_2”.

Once all REPINs in a single genome for a single REP sequence group were identified, we compared the conserved parts of each REPIN (total length of 42bp). Because these data also contain sequences that are likely not to be mobile anymore we defined a REPIN population as all REPINs that differ by no more than three nucleotides from the master sequence. Clusters consisting solely of REP sequences were ignored.

### Identify RAYTs in the genome

To identify RAYT genes in *Neisseria* we ran TBLASTN with the RAYT protein (NMAA_0235) from *N. meningitidis* WUE 2594 on 130 *N. meningitidis* and *N. gonorrhoeae* genomes with an e-value cut-off of 1e-80 (only RAYTs mobilizing REPINs are identified, other RAYTs are ignored). For *P. chlororaphis* we used the RAYT protein (PFLU4255) from *P. fluorescens*

SBW25 on 42 *P. chlororaphis* genome, with an e-value cut-off of 1e-20 (three divergent RAYT sequences associated with each of the REPIN populations can be identified). The datasets can be downloaded under http://doi.org/10.5281/zenodo.5139700 (*P. chlororaphis*) (68) and http://doi.org/10.5281/zenodo.5139705 (*Neisseria*) (65).

RAYT alignments were performed using MUSCLE with default parameters (69). Phylogenies were calculated in PHYML (45).

Whether RAYTs belong to a specific REPIN population (colour) in *P. chlororaphis* was determined in three different ways. First, RAYTs that are associated with a specific REPIN population are also flanked by a REP sequence / REPIN from this population. Second, RAYTs that are associated with the same REPIN population are found in the same extragenic space in the genome (between the same genes). Third, a phylogenetic analysis shows that all RAYTs of the same group form monophyletic clades in the phylogeny (**Supplementary Figure 1**).

### Analyzing 16mer repeats across bacterial genomes

We downloaded complete, representative bacteria from NCBI (http://www.ncbi.nlm.nih.gov/genome/browse/) on the 16.11.2020. This resulted in a list of 2667 genomes (2828 chromosomes). For each of the chromosomes we determined the frequency of the most abundant 16mer.

We also determined whether a genome contained a RAYT gene via TBLASTN (70) using PFLU4255 from SBW25 (e-Value threshold set to 0.01). The amino acid sequences from all identified RAYTs were then compared to each other using the Needleman–Wunsch algorithm. To the distance matrix we applied MCL with standard settings to identify sequence clusters as described previously (see (18) for details). IS*200* genes were identified via TBLASTN using an IS*200* gene from *E. coli* RHB29-C15 gene QMG24152.1 (e-Value threshold set to 0.001). If a match had an e-Value of above 0.001, then the genome was designated an IS*200* containing genome.

### Calculating dN/dS values

We calculated dN/dS using the KaKs Calculator version 1.2 using the Nei Gojobori model (NG) (71, 72). We applied the KaKs calculator to RAYT alignments performed in Geneious 2022.2.2 using the translation align method, which employs MUSCLE to perform the protein alignments (69, 73). Across all pairwise KaKs values we reported the mean in the manuscript.

## Acknowledgements

We thank Jenna Gallie for helpful comments on the manuscript, Eric Hugoson for bioinformatics support, generous core funding from the Max Planck Society, and support from the Deutsche Forschungsgemeinschaft (DFG) Collaborative Research Center 1182 ‘Origin and Function of Metaorganisms’ (grant no. SFB1182) to PBR.

## Supplementary Figures

**Supplementary Figure 1.**
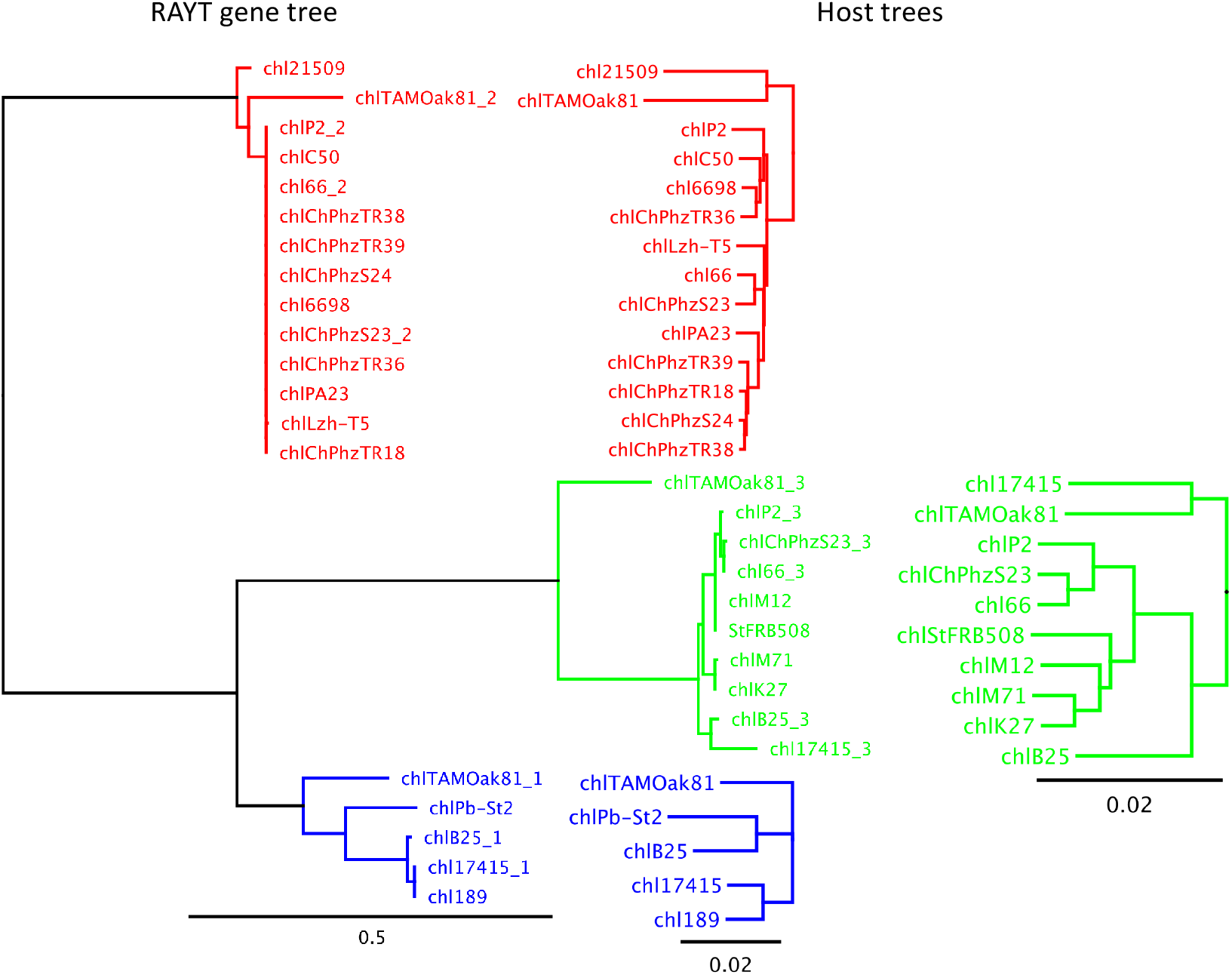
The figure shows the RAYT phylogeny on the left and the corresponding host phylogeny for each of the RAYTs on the right side. The host phylogenies were determined from complete genome sequences using andi (48) and PHYML (45). The RAYT phylogeny was built from alignments generated with MUSCLE (69) and the phylogeny was inferred with PHYML (45).

**Supplementary Figure 2.**
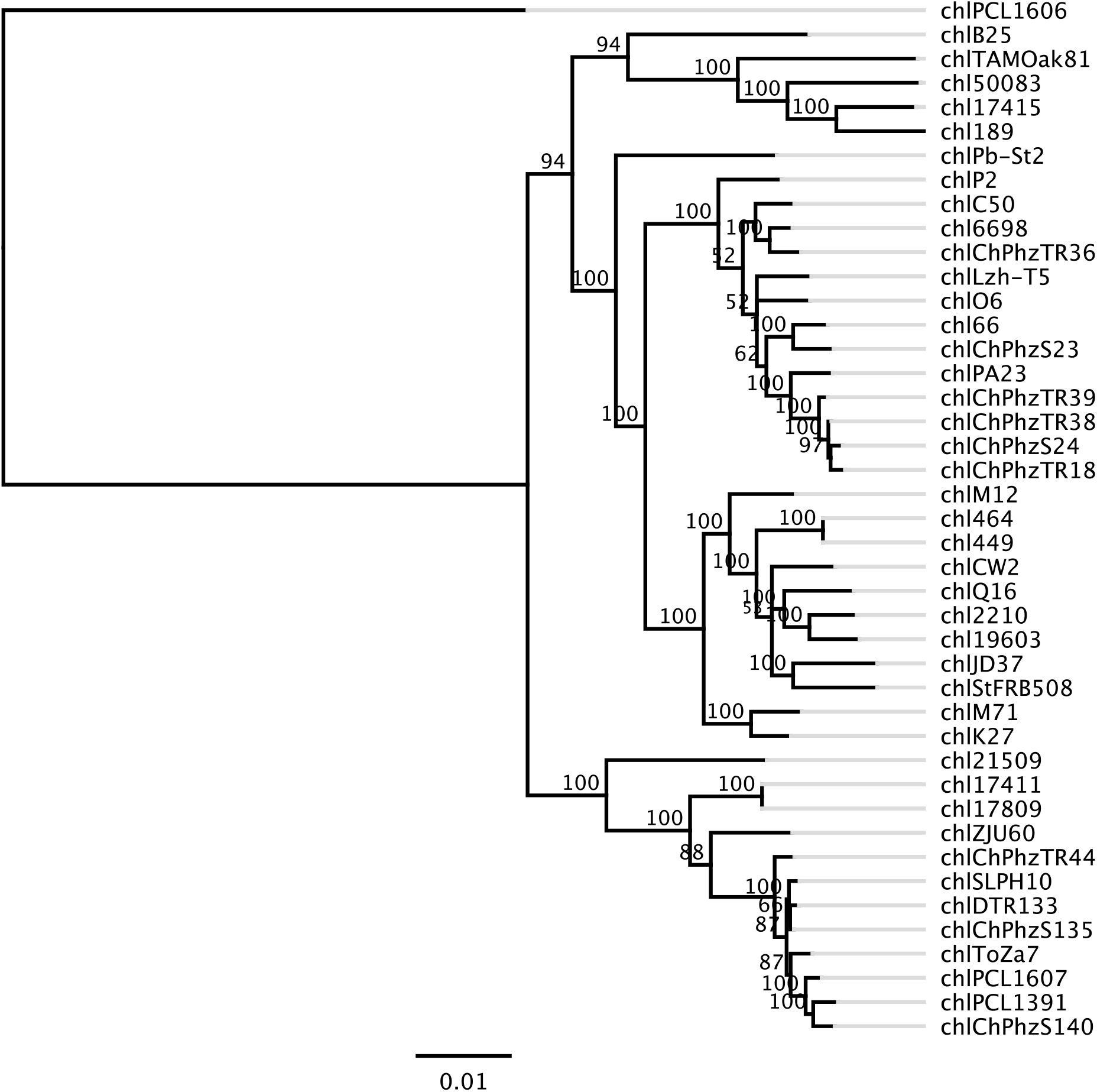
Whole genome phylogeny of all *P. chlororaphis* genomes. The whole genome alignment was generated via REALPHY (44). REALPHY was run on the webserver (version 1.13) using TAMOak and 17411 as references. The resulting alignments were merged resulting in a total of 3,850,960 alignment positions. From this alignment a phylogeny and 100 bootstrap replicates were generated with a GTR substition model using PhyML (45).

**Supplementary Figure 3.**
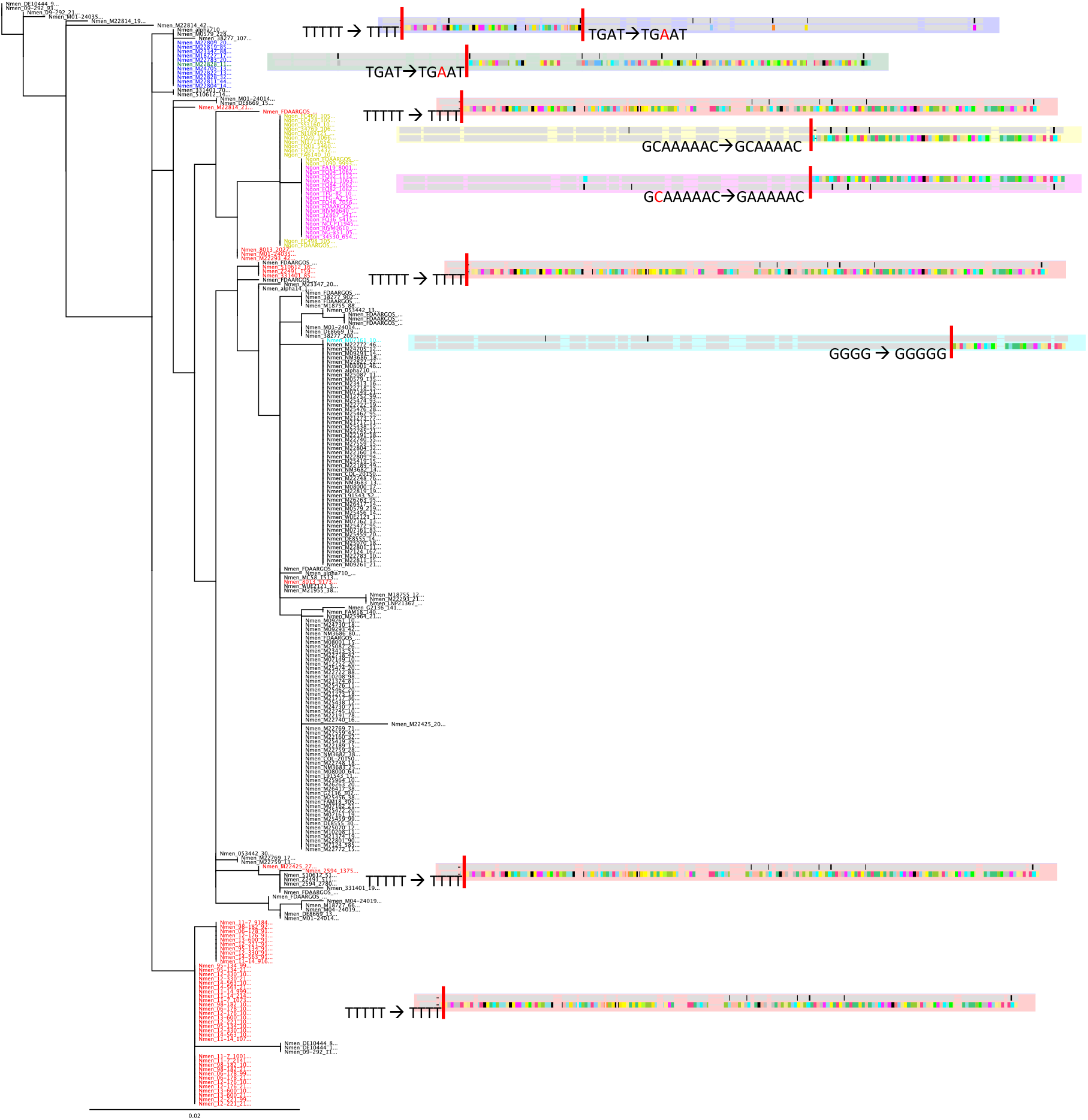
Phylogeny of all *Neisseria* RAYTs. RAYT genes are named by the *Neisseria* strain in which they are contained. All RAYTs from *N. gonorrhoeae* contain a frameshift mutation (yellow and purple). Different colours indicate different frameshift mutations. Note that the red frameshift mutation occurs multiple times independently across the phylogeny and that occasionally frameshift mutations revert. Red bars indicate the position of the frameshift in the RAYT gene. The phylogeny was built from a MUSCLE (69) alignment using PHYML (45).

